# New complexities of SOS-induced “untargeted” mutagenesis in *Escherichia coli* as revealed by mutation accumulation and whole-genome sequencing

**DOI:** 10.1101/2020.01.15.908301

**Authors:** Brittany A. Niccum, Christopher P. Coplen, Heewook Lee, Wazim Mohammed Ismail, Haixu Tang, Patricia L. Foster

**Affiliations:** Department of Biology, Indiana University, Bloomington, IN, USA, 47405; Luddy School of Informatics, Computing and Engineering, Indiana University, Bloomington, IN, USA, 47405

**Keywords:** SOS response, SOS mutagenesis, mutation accumulation, mutation hotspots, DNA polymerase fidelity, error-prone DNA polymerases

## Abstract

When its DNA is damaged, *Escherichia coli* induces the SOS response, which consists of about 40 genes that encode activities to repair or tolerate the damage. Certain alleles of the major SOS-control genes, *recA* and *lexA*, cause constitutive expression of the response, resulting in an increase in spontaneous mutations. These mutations, historically called “untargeted”, have been the subject of many previous studies. Here we re-examine SOS-induced mutagenesis using mutation accumulation followed by whole-genome sequencing (MA/WGS), which allows a detailed picture of the types of mutations induced as well as their sequence-specificity. Our results confirm previous findings that SOS expression specifically induces transversion base-pair substitutions, with rates averaging about 60-fold above wild-type levels. Surprisingly, the rates of G:C to C:G transversions, normally an extremely rare mutation, were induced an average of 160-fold above wild-type levels. The SOS-induced transversion showed strong sequence specificity, the most extreme of which was the G:C to C:G transversions, 60% of which occurred at the middle base of 5′GGC3′+5′GCC3′ sites although these sites represent only 8% of the G:C base pairs in the genome. SOS-induced transversions were also DNA strand biased, occurring, on average, 2- to 4- times more often when the purine was on the leading strand template and the pyrimidine on the lagging strand template than in the opposite orientation. However, the strand bias was also sequence specific, and even of reverse orientation at some sites. By eliminating constraints on the mutations that can be recovered, the MA/WGS protocol revealed new complexities to the nature of SOS “untargeted” mutations.

**Highlights:**

- The SOS response to DNA damage induces “untargeted” mutations
- SOS-mutations are revealed by mutation accumulation and whole genome sequencing
- SOS-mutations are both sequence and DNA-strand biased
- G:C to C:G transversions are particularly highly induced by SOS
- G:C to C:G transversions are extremely sequence and DNA-strand biased

## Introduction

When experiencing DNA damage bacteria induce the SOS response that consists of activities to repair or tolerate the damage. In *E. coli* the SOS response comprises a regulon of about 40 genes that are transcriptionally repressed by the LexA protein. RecA, *E. coli*’s recombinase, induces the response by first interacting with the single-stranded DNA that is produced when DNA is damaged. RecA and DNA form nucleoprotein filaments, historically known as RecA*, that act as coproteases, inducing LexA to self-cleave and, thus, become inactive. LexA inactivation results in the transcriptional induction of the SOS genes. Both the *lexA* and *recA* genes are themselves repressed by LexA, as are the *polB, dinB*, and *umuDC* genes that encode the specialized DNA polymerases, Pol II, Pol IV, and Pol V, respectively. Lacking proofreading functions, Pol IV and Pol V are error-prone, whereas Pol II is accurate. RecA* also induces the self-cleavage of the UmuD protein, a dimer of which forms a complex with UmuC to make the active Pol V polymerase. But in order to synthesize DNA, Pol V must be stimulated by interacting with RecA*. Pol V is responsible for most of the mutagenic consequences of DNA damage in *E. coli*. (for a review of the SOS response, see [1].

In the absence of overt DNA damage, the SOS response can be induced by genetically modifying the *recA* gene to produce constitutive RecA* activity. Inactivating the *lexA* gene further stimulates the response. Both modifications can only be accomplished in a strain that is defective for *sulA* (formerly known as *sfiA*), an SOS gene that encodes an inhibitor of septation. In a *sulA*^+^ strain, constitutive expression of the SOS response results in lethal filamentation [2].

Constitutive expression of the SOS response is mutagenic, mainly due to the activity of Pol V [3-5], and, to a lesser extent Pol IV [6, 7]. Other DNA polymerases also play a role under certain circumstances [8]. The nature of these mutations has been investigated for many years. Originally named “untargeted” mutagenesis [9], to distinguish the process from the SOS-dependent mutagenesis caused by DNA damage from exogenous agents, it has been debated whether the resulting mutations are truly untargeted or are due to spontaneous endogenous DNA damage. Alternatively, the increased mutation rate could result from Pol V or Pol IV being able to access the replication fork because of their increased cellular concentration when SOS is constitutive. The polymerases could then synthesize a track of error-containing DNA, or, because they lack proofreading activity, extend mismatches made by the replicase, Pol III, that would otherwise be corrected [10, 11].

To try to distinguish among these hypotheses, several groups have studied the specific mutations induced by constitutive expression of the SOS response. In general, SOS mutations differ from the mutations induced by most types of DNA damage, arguing against the hypothesis that the mutations are due to endogenous lesions. The roles of the various polymerases have also been deduced by comparing the types of SOS mutations to the errors that the polymerases are known to make [8, 11]. However, these studies invariably used mutation detection assays that rely on phenotypic selection, which means only certain mutations can be detected. Even *in vitro* assays of polymerase fidelity rely on a specific site or stretch of DNA as template. Inevitably these restrictions limit and bias the types of mutations that can be recovered.

In the study reported here we utilized mutation accumulation followed by whole genome sequencing to revisit the nature of the SOS “untargeted” mutations. In the mutation accumulation protocol, strains of bacteria are put through repeated single-cell bottlenecks, a procedure that reduces the selective pressure against most mutations except for very deleterious ones. This protocol allows for the accumulation of a population of nearly neutral mutations. Whole genome sequencing of the resulting bacteria recovers mutations at all possible sequence targets, removing the bias due to sequence constraints. The results we obtained confirm many previous findings, but also offer some new information about the nature of SOS untargeted mutations.

## Materials and Methods

### Bacterial strains

The strains used in this study are listed in Table S1 in Appendix A and are derivatives of PFM2 [12]. Strains were made Δ*sulA* by P1 transduction [13] of Δ*sulA*::Kn^R^ from strain JW0941 [14] into the recipient, selecting for Kn^R^, followed by FLP recombination to remove the Kn^R^ element [15]. Strains were made *recA730* [16] by first creating a Δ*recA*::*cat*-ISceI cassette and crossing it onto the chromosome as described [17]. The Δ*recA*::*cat*-ISceI allele was replaced with *recA730* by transformation with the PCR product of the *recA730* mutant strain, PC1427 [18] selecting for resistance to anhydrotetracycline, which indicates that the ISceI cut sites are eliminated. The *lexA3* allele [19] was moved into strains by P1 transduction of Δ*metA*::Kn^R^ from strain JW3973 [14] into the recipient, followed by P1 transduction to *metA*^+^ *lexA3* from strain FC231 [20], selecting for growth on minimal medium. To make a *lexA* deletion allele, we used PCR with pKD13 [15] to create a cassette consisting of the *lexA* promoter and the first 8 Nt of *lexA*, the Kn^R^-element surrounded by FRT sites, and the *lexA* sequence from Nt 479 to 530. The cassette was then recombined into PFM117, a Δ*sulA* derivative of PFM2 carrying the λ phage Red recombinase expression plasmid pKD46 [15]. After heat-elimination of pKD46, the *lexA*-Kn^R^ cassette was moved by P1 transduction into appropriate strains and then the *lexA*-Kn^R^ cassette was removed by FLP recombination, as described [15]. This created a deletion of *lexA* from Nt 9 to Nt 478, leaving an 82 Nt in-frame scar that should not be polar on the downstream *dinF* gene. Construction of Δ*umuDC* and Δ*dinB* mutant strains has been previously described [21]. After construction, all strains were confirmed by appropriate phenotypes and sequencing. The Δ*lexA* allele was also confirmed by Western blotting (R. Woodgate and J.P. McDonald, personal communication).

### Media

Strains were grown in Miller Luria Broth (LB) (Difco; BD) and streaked on the same medium plus 15 g/L BactoAgar (Difco). When required, antibiotic concentrations were: carbenicillin (Carb), 100 μg/ml; kanamycin (Kn), 50 μg/ml; nalidixic acid (Nal), 40 μg/ml; chloramphenicol (Cam), 30 μg/ml; rifampicin (Rif), 100 μg/ml; and, anhydrotetracycline (AHT), 500 μg/mL.

### Mutation accumulation experiments

The parameters of the mutation accumulation (MA) experiments described in this paper are given in Table S2 in Appendix A. The MA protocol has been described in our previous papers [12, 21, 22]. Multiple parallel MA lines were initiated from one or two founder colonies, as described [22]. Every day each MA line was streaked for single colonies on LB-agar plates that were then incubated at 37°C. The number of passes required to achieve a statistically meaningful number of mutations was estimated from preliminary fluctuation assays, as described [12, 21, 22]. At the end of the MA experiment, a single colony of each line was grown in LB broth overnight and a freezer stock made.

### Mutation rates

As described [12, 21, 22], the mutation rate for each MA experiment was calculated by dividing the total number of mutations accumulated by the total number of generations undergone by all the MA lines. For the conditional mutation rates, this value was then divided by the appropriate number of sites (A:T, G:C, etc.). The mutation rates for each MA line in an experiment was used to compute the variance for statistical analysis, as described [22].

### Estimation of generations

At each passage, the diameter of the colony used for streaking was measured. The number of CFUs in colonies of appropriate diameters was then determined and used to estimate the number of generations that occurred in each MA line, as described [12, 21, 22]

### Genomic DNA preparation. library construction, and sequencing

PureLink Genomic DNA purification kits (Invitrogen) were used to extract genomic DNA from overnight LB-broth cultures inoculated from the freezer stocks. Paired end libraries were prepared by the Indiana University Center for Genomics and Bioinformatics and were sequenced by the University of New Hampshire Hubbard Center for Genome Studies using the Ilumina HiSeq 2500 platform. Insert size was between 350 and 600 base pairs.

### Sequence analysis

The source code used for SNP calling is entitled “MA-pipeline” and is available at https://github.com/COL-IU/MA-pipeline. The *E. coli* reference genome NCBI NC_000913.2 was used. Illumina reads were aligned to the reference genome using the Burrows-Wheeler short-read alignment tool, BWA version 0.7.9 [23]. A MA line would be eliminated if its sequence coverage was poor or if it shared more than 50% of its mutations with another line (because of mutations occurring in the founder colony, or cross contamination during streaking). If fewer than 50% of the mutations were shared, mutations were assigned to each line at random. The sequences and SNPs reported in this paper have been deposited in the National Center for Biotechnology Information Sequence Read Archive https://trace.ncbi.nlm.nih.gov/Traces/sra/ (accession no. PRJNA168337) and in the IUScholarWorks Repository (hdl.handle.net/to be determined).

### Determination of DNA strand bias

Strand bias was determined as described [22]. Basically, because of *E. coli*’s bidirectional mode of DNA replication, the chromosome is divided into two independently replicating halves, called replichores. A triplet read 5′ to 3′ in the reference sequence is on the lagging-strand template (LGST) on the right replichore and on the leading-strand template (LDST) on the left replichore. The complement to the triplet has the opposite orientation.

## Results

### Constitutive expression of the SOS regulon increases mutation rates

The RecA730 protein contains an E38K mutation that makes it constitutive for all SOS functions including LexA and UmuD cleavage [24]. The *recA730* allele is lethal unless the *sulA* gene is inactive, and *sulA* is deleted in all the *recA730* mutant strains reported here. A mutant accumulation/whole genome sequencing (MA/WGS) experiment with a *recA730* mutant strain revealed a 30-fold increase in the rate of base-pair substitutions (BPSs) (Table 1) and a 13-fold increase in the rate of small indels (≤ 4 bp) relative to wild-type rates (Table 2). These increases are in line with those obtained in other studies with SOS constitutive mutant strains [4, 5, 25-27]. When the *recA730* mutant strain also carried a deletion of most of the *lexA* gene (see Materials and Methods), the BPS rates increased another 40% (Table 1), reflecting the fact that LexA is not completely inactivated in *recA730* mutant strains (R. Woodgate and J.P. McDonald, personal communication). In contrast, the rate of indels declined by about 40% when *lexA* was absent (Table 2), but this difference is based on very small numbers (28 vs 16 indels).

**Table 1:**
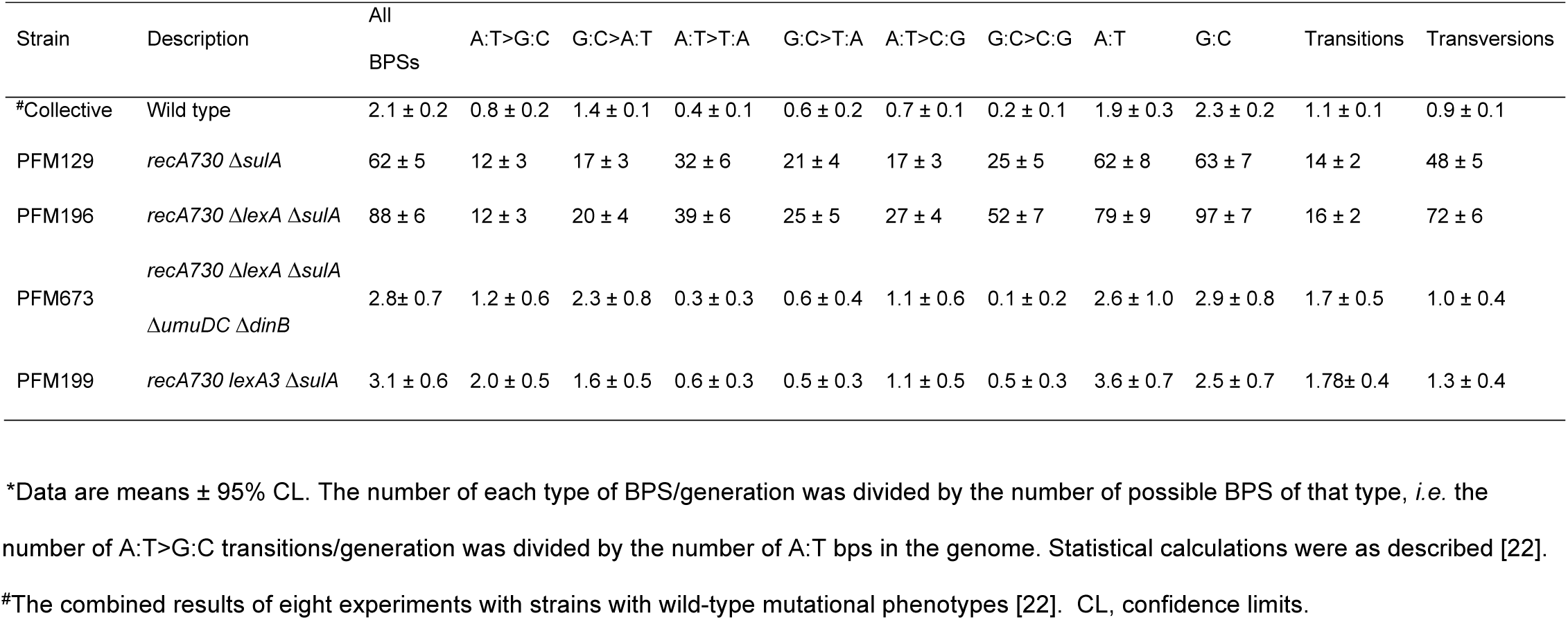
*SOS-induced conditional BPS rates × 10^10^.

| Strain | Description | All<br>BPSs | A:T>G:C | G:C>A:T | A:T>T:A | G:C>T:A | A:T>C:G | G:C>C:G | A:T | G:C | Transitions | Transversions |
| --- | --- | --- | --- | --- | --- | --- | --- | --- | --- | --- | --- | --- |
| #Collective | Wild type | 2.1 ± 0.2 | 0.8 ± 0.2 | 1.4 ± 0.1 | 0.4 ± 0.1 | 0.6 ± 0.2 | 0.7 ± 0.1 | 0.2 ± 0.1 | 1.9 ± 0.3 | 2.3 ± 0.2 | 1.1 ± 0.1 | 0.9 ± 0.1 |
| PFM129 | *recA730 ΔsulA* | 62 ± 5 | 12 ± 3 | 17 ± 3 | 32 ± 6 | 21 ± 4 | 17 ± 3 | 25 ± 5 | 62 ± 8 | 63 ± 7 | 14 ± 2 | 48 ± 5 |
| PFM196 | *recA730 ΔlexA ΔsulA* | 88 ± 6 | 12 ± 3 | 20 ± 4 | 39 ± 6 | 25 ± 5 | 27 ± 4 | 52 ± 7 | 79 ± 9 | 97 ± 7 | 16 ± 2 | 72 ± 6 |
| PFM673 | *recA730 ΔlexA ΔsulA*<br>*ΔumuDC ΔdinB* | 2.8 ± 0.7 | 1.2 ± 0.6 | 2.3 ± 0.8 | 0.3 ± 0.3 | 0.6 ± 0.4 | 1.1 ± 0.6 | 0.1 ± 0.2 | 2.6 ± 1.0 | 2.9 ± 0.8 | 1.7 ± 0.5 | 1.0 ± 0.4 |
| PFM199 | *recA730 lexA3 ΔsulA* | 3.1 ± 0.6 | 2.0 ± 0.5 | 1.6 ± 0.5 | 0.6 ± 0.3 | 0.5 ± 0.3 | 1.1 ± 0.5 | 0.5 ± 0.3 | 3.6 ± 0.7 | 2.5 ± 0.7 | 1.78 ± 0.4 | 1.3 ± 0.4 |

\*Data are means ± 95% CL. The number of each type of BPS/generation was divided by the number of possible BPS of that type, *i.e.* the number of A:T>G:C transitions/generation was divided by the number of A:T bps in the genome. Statistical calculations were as described [22].
#The combined results of eight experiments with strains with wild-type mutational phenotypes [22]. CL, confidence limits.

**Table 2:** *SOS-induced conditional Indel rates × 10^10^.

| Strain | Description | Total | +1A:T | -1A:T | +1G:C | -1G:C | +>1 bp | ->1 bp | In run >2 bp |
| --- | --- | --- | --- | --- | --- | --- | --- | --- | --- |
| #Collective | Wild type | 0.21 ± 0.04 | 0.07 ± 0.03 | 0.19 ± 0.06 | 0.03 ± 0.01 | 0.10 ± 0.07 | 0.002 ± 0.003 | 0.01 ± 0.01 | 1.01 ± 0.28 |
| PFM129 | *recA730 ΔsulA* | 2.9 ± 0.9 | 0.60 ± 0.69 | 2.4 ± 1.4 | < 0.2 | 2.1 ± 1.3 | 0.10 ± 0.20 | 0.10 ± 0.21 | 8.3 ± 4.2 |
| PFM196 | *recA730 ΔlexA ΔsulA* | 1.8 ± 1.2 | 0.44 ± 0.66 | 1.6 ± 2.0 | 0.22 ± 0.44 | 1.3 ± 1.0 | < 0.1 | < 0.1 | 3.3 ± 2.9 |
| PFM673 | *recA730 ΔlexA ΔsulA ΔumuDC ΔdinB* | 0.37 ± 0.23 | < 0.08 | 0.59 ± 0.42 | 0.08 ± 0.17 | < 0.08 | < 0.04 | 0.04 ± 0.08 | 1.5 ± 1.2 |
| PFM199 | *recA730 lexA3 ΔsulA* | 0.32 ± 0.19 | 0.09 ± 0.12 | 0.21 ± 0.18 | 0.04 ± 0.09 | 0.21 ± 0.24 | 0.04 ± 0.06 | < 0.02 | 1.40 ± 0.92 |

\*Data are means ± 95% CL. The number of each type of indel/generation was divided by the number of possible indels of that type, *i.e.* the number of +1A:T insertions/generation was divided by the number of A:T bps in the genome; for indels involving more than one bp, the indel/generation was divided by the number of bp in the entire genome; for indels at runs greater than 2 bp, the number of indels/generation at runs >2 bp was divided by the number of bps in the genome in such runs, which is 765441. Statistical calculations were as described [22]
#The combined results of eight experiments with strains with wild-type mutational phenotypes [22]. CL, confidence limits

When the *recA730* mutant strain carried the *lexA3* allele that encodes a non-cleavable LexA protein [28], the mutation rate dropped to near wild-type levels (Tables 1 & 2). Eliminating the error-prone polymerases Pol IV and Pol V by deleting the *dinB* and *umuDC* genes likewise reduced the mutation rate of the *recA730* Δ*lexA* mutant strain to near wild-type levels (Tables 1 & 2). In our experiments we did not distinguish between the roles of Pol IV and Pol V, but, as mentioned above, previous reports have shown that SOS mutagenesis is strongly dependent on Pol V [3-5] and that Pol IV may also play a role [6-8]. *Transversion mutations are highly induced when SOS is constitutive*

It has been known for some time that constitutive expression of the SOS regulon not only stimulates transition BPSs, but especially stimulates transversion BPSs [5, 25-27]. The results of our MA/WGS results confirm these prior results and reveal additional mutagenic specificity (Figure 1). In the SOS constitutive strains, the rates of transition mutations were increased 14- to 15-fold over wild-type rates, and G:C to A:T and A:T to G:C mutations were stimulated about equally. The rates of transversion BPSs were increased 50- to 80-fold overall, but rates varied depending on the mutagenic event. Compared to wild-type rates, G:C to T:A and A:T to C:G BPSs were stimulated 25- to 40-fold while A:T to T:A BPSs were stimulated 100-fold. Surprisingly, the rate of G:C to C:G BPSs, normally a rare mutation, was stimulated 100- to 200-fold relative to the wild-type levels. These results will be compared to previous findings in the Discussion section.

**Figure 1.**
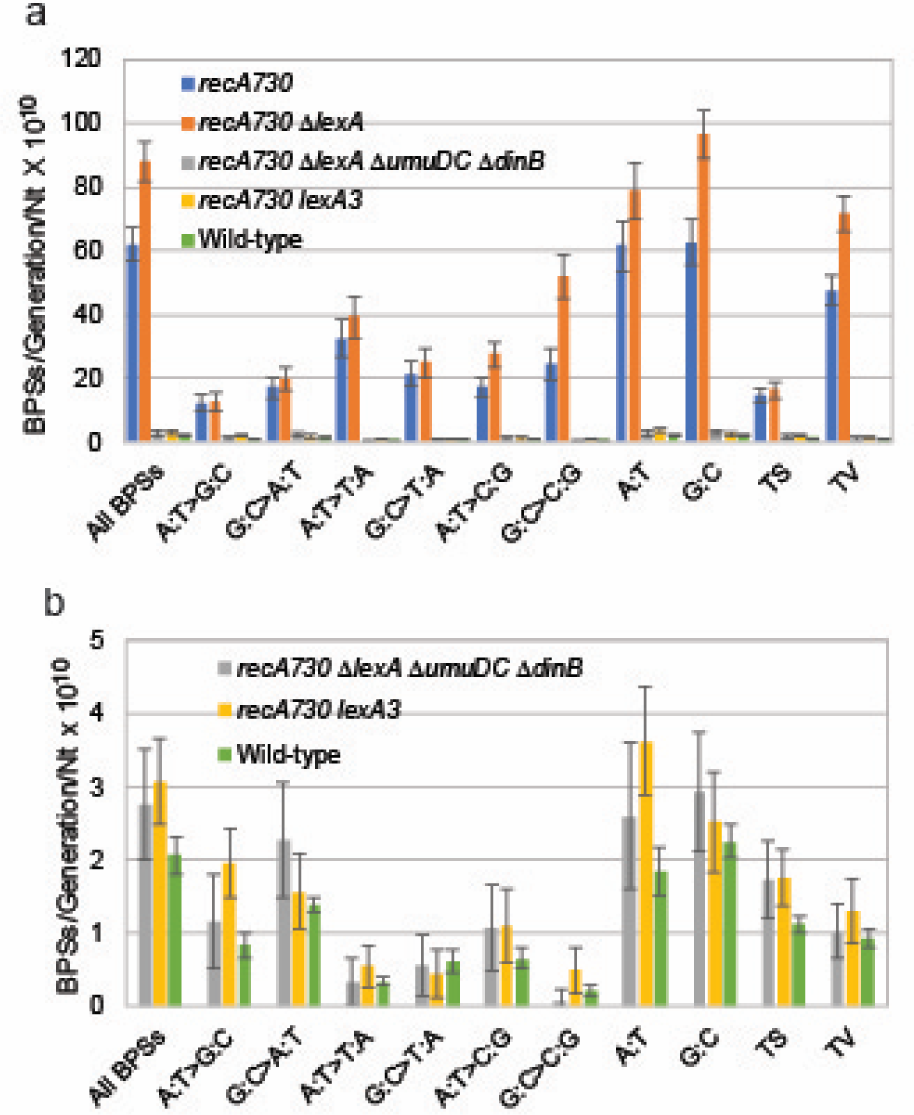
Base-pair substitutions induced by constitutive expression of the SOS response. Figure 1a. The conditional mutation rates of SOS-induced base pair substitutions and their dependence on DNA Pol V and Pol IV, encoded by the *umuDC* and *dinB* genes, respectively, and repression by the non-inducible LexA protein encoded by the *lexA3* allele. Conditional mutation rates are the rate per generation divided by the number of Nt of the relevant type in the genome. Figure 1b. As Fig. 1a, but with the data for the SOS-constitutive strains, *recA730* and *recA730* Δ*lexA*, removed to allow comparisons of the data from wild-type strains to the SOS-inactive strains, *recA730* Δ*umuDC* Δ*dinB* and *recA730 lexA3*. Bars represent means and the error bars are 95% CLs computed as described [22]. TS, transitions; TV, transversions. *recA730*, PFM129; *recA730* Δ*lexA*, PFM196; *recA730* Δ*lexA* Δ*umuDC* Δ*dinB*, PFM673; *recA730 lexA3*, PFM199; Wild-type, eight MA experiments combined [22]. All r*ecA730* strains are also Δ*sulA*. CL, confidence limits.

### SOS BPSs rates are site-specific

The different types of BPSs induced by constitutive expression of the SOS regulon are not randomly distributed but are concentrated at certain DNA sequences. The bps immediately 3’ and 5’ to a bp appear to determine its mutability (Figure 2); we have been unable to identify longer sequence motifs that might affect mutation rates. As shown in Figure 2, in general the difference in mutation rate between the most and least mutable sites was about 10-fold for transitions and 20-fold for transversions. Of particular note, the rate of G:C to C:G transversions at 5′G**G**C3′+5′G**C**C3′ sites was 40-fold greater than the rate of this transversion at the least mutable site,5′C**G**A3′+5′T**C**G3′ (Figure 2f) (throughout this paper, a triplet and its complement are presented 5′ to 3′ with the mutated base in the center and in bold). Indeed, of the 368 G:C to C:G transversions recovered, 222 (60%) were at 5′G**G**C3′+5′G**C**C3′ sites, yet these sites represent only 8% of the G:C bps in the genome.

**Figure 2.**
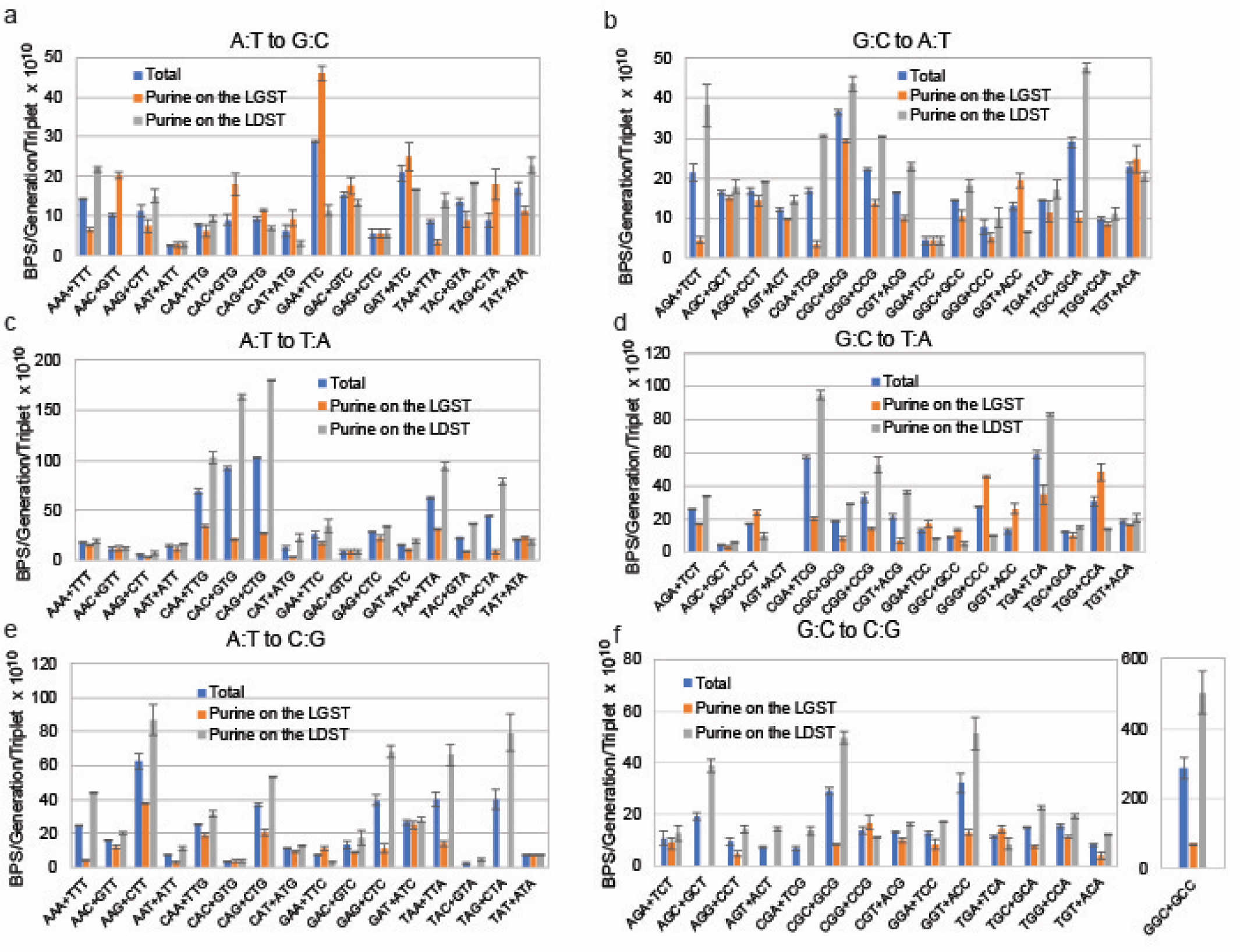
The site and strand specificity of the SOS-induced base-pair substitutions. Conditional mutation rates are computed from the combined results of the *recA730* (PFM129) and *recA730* Δ*lexA* (PFM196) mutant strain both of which are also Δ*sulA*. LGST = DNA lagging strand template; LDST = DNA leading strand template. Triplets are written 5′ to 3′ with the target base in the center. Bars represent means and the error bars are 95% CLs computed as described [22]. Note change of scales among the charts and the extra chart in Fig. 2f for results at 5′G**G**C3′ + 5′G**C**C3′ sites. CL, confidence limits.

### SOS BPS are DNA-strand biased

In a previous report [29], the DNA-strand biases of three mutations (G:C to T:A, A:T to T:A, and A:T to G:C) in the *lacZ* gene were assayed in a *recA730 lexA*^+^ *mutL* (mismatch repair defective) mutant strain. Maliszewska-Tkaczyk *et al*. (2000) found that the frequencies of G:C to T:A and A:T to T:A BPSs were 2 and 12-fold higher, respectively, when the purine was on the leading-strand template (LDST) and the pyrimidine was on the lagging-strand template (LGST) than in the opposite orientation. We partially confirmed this result (Figure 2, Figure S1 in Appendix A); for all four transversion mutations, the overall rate of SOS-induced BPSs was about 3-fold higher when the purine was on the LDST rather than on the LGST. However, the bias varied among mutation types. The rate of G:C to C:G transversions was nearly 5-fold higher when the G was on the LDST rather than on the LGST, whereas that ratio was only 1.5-fold for G:C to T:A transversions (Figure S1 in Appendix A). The strand bias also varied among sequence contexts; for example, the rate of G:C to T:A transversions with G on the LDST compared to on the LGST was 5-fold higher at 5′C**G**T3′+5′A**C**G3′ sites, but was almost 5-fold lower at 5′G**G**G3′+5′C**C**C3′sites, even though the overall rate of G:C to T:A BPSs was nearly the same at the two sites (Figure 2d, Table S3 in Appendix A). (Throughout this paper triplet sequences are given 5′ to 3′ with the mutated base in the center).

In contrast to the transversion mutations, our results showed that A:T to G:C transitions had a slight (1.3x) bias for rates to be higher when the A was on the LGST. However, here, too, that were large differences among sites (Figure 2a, Table S3 in Appendix A). At 5′A**A**A3′+5′T**T**T3′ sites, the rate of A:T to G:C transitions was 3-fold higher when the A was on the LDST compared to on the LGST, whereas at 5′G**A**A3′+5′T**T**C3′ sites, the bias was reversed, with the rate 5-fold higher when the A was on the LGST rather than on the LDST. G:C to A:T transitions were biased for the G to be on the LDST at almost all sites (Figure 2b, Table S3 in Appendix A). The single exception with statistical significance was at 5′G**G**T3′+5′A**C**C3′ sites, where the rate of G:C to A:T transitions was 3-fold higher when the G was on the LGST rather than on the LDST (P = 0.03).

### The spectrum of SOS-induced indels resembles that of wild-type

Although constitutive expression of the SOS regulon increased the rate of all types of indels about 13-fold relative to wild-type rates, the spectrum of indels remained about the same (Figure 3; Table 2). For both SOS-constitutive and wild-type strains, loss of a bp was more frequent than gain of one, loss of A:Ts and G:Cs occurred at about the same rate, and runs were hotspots. It appears from Figure 3b that indels at runs were relatively less frequent in the SOS strains than in the wild-type strains, but the numbers are small, and there was a large, unexplained, difference in the number of indels in runs between the *recA730 lexA*^+^ strain and the *recA730* Δ*lexA* strain (14 vs 5). This difference largely accounts for the lower rate of indels observed in the *recA730* Δ*lexA* strain compared to the *recA730 lexA*^+^ strain (Table 2).

**Figure 3.**
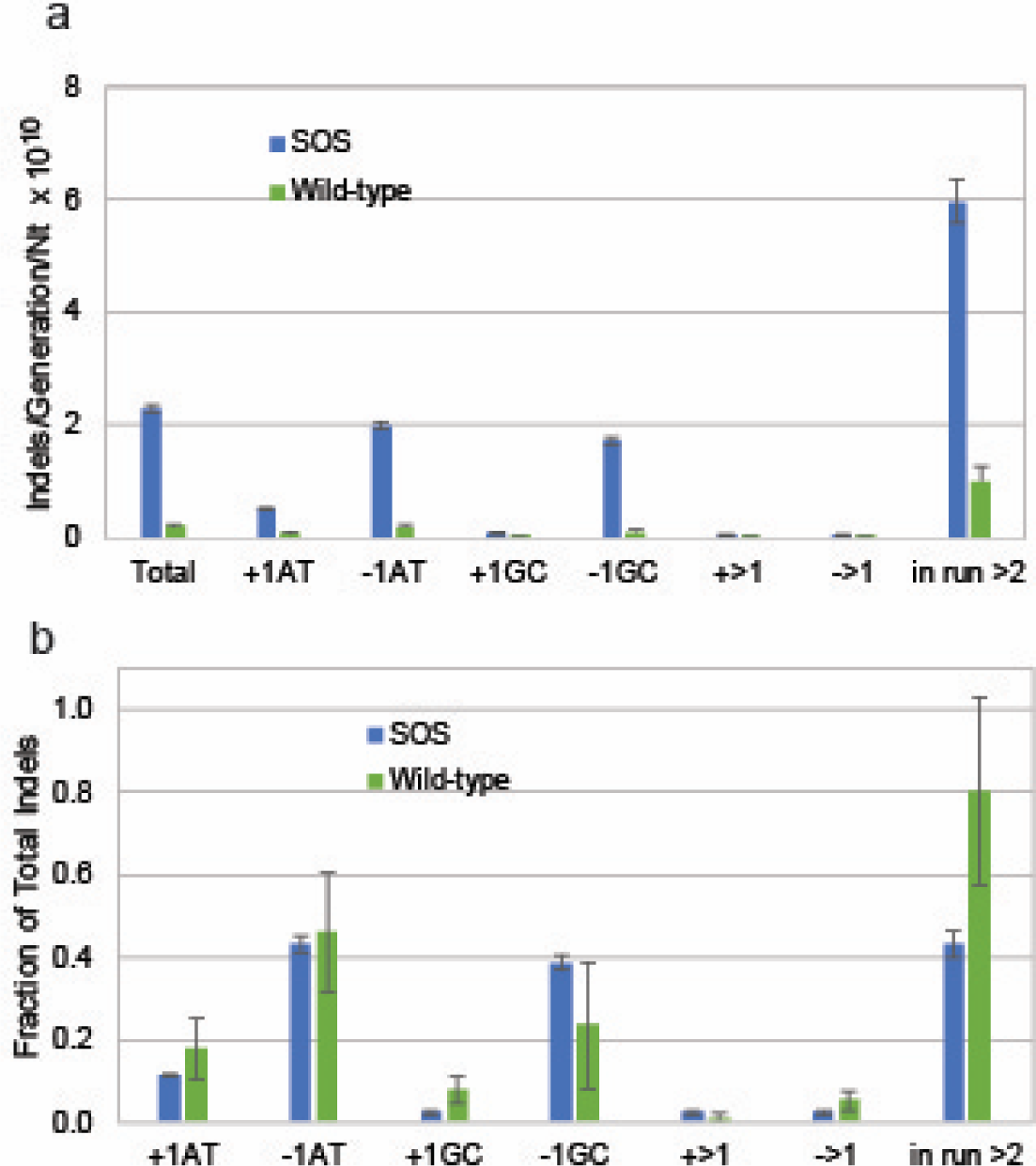
Indels induced by constitutive expression of the SOS response. Fig. 3a. The conditional mutation rates of the SOS-induced indels. Because of the low numbers of indels induced, the results of the *recA730* and *recA730* Δ*lexA* mutant strains have been combined. For the rates of indels >1 Nt, the rate per generation was divided by the total number of Nt in the genome; for the rates of indels in runs, the rate per generation was divided by the number of Nt in runs; for all other types of indels, the rate per generation was divided by the number of Nt of the relevant type in the genome. Fig. 3b. The fractions of each type of indel comparing the SOS-induced results with the results from the wild-type strains. The fractions were calculated using the mutation rates per generation. Bars represent means and the error bars are 95% CLs computed as described [22]; fractions and error bars of the fractions were calculated as described [30]. *recA730*, PFM129; *recA730* Δ*lexA*, PFM196; Wild-type, eight MA experiments combined [22]. All r*ecA730* strains are also Δ*sulA*. CL, confidence limits.

## Discussion

Although the rates and spectra of the mutations induced by constitutive expression of the SOS response have been studied previously, we undertook our MA experiments to explore the phenomenon across the entire genome. This allowed us to analyze the influence of many different sequence contexts on the types and rates of the induced mutations. We confirmed previous findings [5, 25-27] that transversion mutations are more highly induced by SOS expression than are transition mutations (Figure 1, Table 1). We further found that that the rates of indels were increased about the same degree as transition mutations (Figure 3, Table 2). In addition, our results reveal that in SOS induced strains G:C to C:G transversions, normally a rare mutation, occur at rates equivalent to or higher than the rates of the other transversions (Figure 1, Table 1), a result that, to our knowledge, has been observed only once before [26].

### The impact of mismatch repair

Several of the studies cited above used strains that were not only expressing the SOS regulon, but that were also defective for mismatch repair (MMR) [7, 8, 27, 29]. We choose not to use such double mutants because the high mutation rate of MMR-defective strains could obscure the relatively milder SOS mutator phenotype. But, mismatch repair imprints its own biases on the mutational spectrum. The efficiency at which MMR prevents BPSs varies with the type of mutation and the local sequence context but is many times greater for transitions than transversions [22]. MMR activity lowers the rate of A:T transitions about 400-fold, and that of G:C transitions about 80-fold, but for transversions these decreases drop to 10-fold or less. The increase in the rates of transversions relative to the wild-type strains in the SOS constitutive strains is 5 to 20-fold greater than the increase due to loss of MMR. Therefore, the spectrum and site specificity of transversions are not complicated by the biases of MMR error-correction.

MMR correction probably determines the spectrum of SOS-induced indels. MMR reduces the rates of most types of indels 100- to 200-fold, but is over 1000-fold efficient at preventing gain of one G:C bp [31], and the rate of this event is particularly low in the SOS constitutive strains (Table 2, Figure 3a). In no case does the indel rate in the SOS constitutive strains exceed that produced by loss of MMR; therefore, MMR is likely active in preventing these mutations and produces a spectrum of SOS-induced indels that resembles that of wild-type strains (Figure 3b). In the discussion below we will focus on the BPSs, particularly transversions.

### Comparison of the results presented here to previous studies

The site-specificities of the SOS-induced BPSs explain, to a large part, the differences between our results and those of previous studies. To our knowledge, all previous investigations of SOS-mutagenesis *in vivo* have relied on mutational assays using phenotypic selection. This inevitably limits the types of mutations that are possible to detect and the sites at which they can occur. In the first of these investigations, Miller and Low [25] identified SOS-induced BPSs by the specific nonsense codons that they created in the *lacI* gene. Because only certain codons can mutate to nonsense codons, the sequence contexts of the mutations were severely limited. Nonetheless, the nonsense results showed that A:T to T:A and G:C to T:A transversions were highly induced by SOS expression, and that G:C to T:A transversions were more frequent when a pyrimidine was 5’ to the G. We confirmed both or these results (Figure 1, Figure 2d). However, Miller and Low (1984) did not detect any A:T to C:G or G:C to C:G transversions. All of the possible sites in the *lacI* gene at which an A:T to C:G transversion can create a nonsense codon have a T 5’ to the target A, which is a disfavored sequence for this mutation (Figure 2e). G:C to C:G transversions can produce a nonsense codon at only 3 sites in the *lacI* gene, and none of these sites have sequences that our data indicate are favored for this mutation (Figure 2f).

Several studies of SOS-mutagenesis used a set of Lac^−^ strains, known as the Cupples strains, each of which revert to Lac^+^ by one of the six possible BPSs [5, 27, 29, 32]. Assaying all six mutations, Watanabe-Akanuma *et al*. (1997) found that the frequencies of A:T to T:A, G:C to T:A, and A:T to C:G transversions were increased in a SOS-constitutive strain 20- to 30-fold relative to wild-type frequencies. In the Lac^−^ strains the triplets containing the target bps for these transversions are 5′C**A**C3′+5′G**T**G3′, 5′C**G**C3′+5′G**C**G3′, and 5′T**A**C3′+5′G**T**A3′, respectively. According to our results (Figure 2 c, d, & e), the first two transversions should be well induced at those sites, but the A:T to C:G transversion should not. Watanabe-Akanuma et al (1997) recovered no G:C to C:G transversions. The sequence context of this mutation In the Lac^−^ strain is 5′T**G**C3′+5′G**C**A3′, which is unfavorable for G:C to C:G BPSs (Figure 2f). Other studies with these same strains will be discussed below.

Rifampicin resistance (Rif^R^) is conferred by 85 BPSs at 60 sites in the *rpoB* gene [33, 34]. Curti *et a*l. [8] used this assay in a *recA730 lexA*(Def) *mutL* strain to analyze the spectrum of BPS produced by the three SOS-induced DNA polymerases, Pol II, IV, and V. They found an overall increase in all transversion mutations except G:C to C:G. While the occurrence of each of these transversions was site-specific, we could find little correlation between the hotspots identified by Curti *et al*. (2009) and the favored triplets identified in our experiments. Of particular note, they found few G:C to C:G mutations despite the fact that there were eight sites for this event, including one with the sequence 5′G**G**C3′+5′G**C**C3′, the dominant triplet in our G:C to C:G data set (Figure 2f). In contrast, in a study of the roles of *E. coli*’s Pol II, IV, and V in spontaneous (not SOS-induced) mutation to Rif^R^, Corzett *et al*. [35] found this site was a modest hotspot for G:C to C:G mutations in wild-type cells and in a strain carrying only Pol V. Understanding these difference results among studies will require further experimentation and analyses.

As mention above, to our knowledge only one study found G:C to C:G transversions to be highly SOS-induced. Yatagai *et al*. [26] sequenced a set of LacI^−^ mutations that were induced by constitutive expression of the SOS regulon. The *lacI* gene is 1082 bp long, and several hundred BPSs will give the LacI^−^ phenotype [36]; thus, this gene presents a large target for mutagenesis. Unfortunately, the majority of LacI^−^ mutations are due to one frameshift hotspot, and so sequencing across the entire genome for BPSs giving the LacI^−^ phenotype is laborious. Yatagai *et al*. (1991) recovered only 42 BPSs, 35 of which were transversions, and 11 of which were G:C to C:G transversions. Seven of the G:C to C:G transversions were at four 5′G**G**C3′+5′G**C**C3′ sites, the sequence that was most favorable for G:C to C:G transversions in our results (Figure 2f). In addition, five G:C to T:A transversions giving the LacI^−^ phenotype occurred at three 5′C**G**A3′+5′T**C**G3′ sites, which was also favorable for G:C to T:A transversions in our data (Figure 2d). Thus, there is a remarkable congruence between our results and those obtained almost 30 years ago.

Indeed, even longer ago the triplet 5′G**G**C3′+5′G**C**C3′ was identified as a transition hotspot in the rII gene of bacteriophage T4 [37]. The transition yielded the nonsense codon TGA and it is not clear if the assay used could have scored transversions.

### Site-specificity of the SOS-induced BPS

The SOS-induced BPS have strong site specificity but is not clear why. As shown in Figure 2, there do not appear to be any evident rules. A:T to T:A transversions are potentiated by a C 5′to the target A (but not at 5′C**A**T3′+5′A**T**G3′), but neither A:T to C:G nor A:T to G:C BPSs share this feature. Instead, A:T to C:G transversions are favored at sites with a G 3′ to the target A, whereas A:T to G:C transitions are favored at sites with a G 5′ to the target A. The strongest sites for G:C to T:A transversions have a pyrimidine 5′ to the G, but only if an A is also 3′ to the G. The most extreme site specificity is G:C to C:G transversions at 5′G**G**C3′+5′G**C**C3′ sites, and, to a much lesser degrees, 5′C**G**C3′+5′G**C**G3′, 5′G**G**G3′+5′C**C**C3′, and 5′G**G**T3′+5′A**C**C3′ sites. But 5′G**G**A3′+5′T**C**C3′ is not a favored site for this mutation. *“dNTP-stabilized misalignment” may contribute to the site specificity of G:C to C:G transversions*

One mechanism that could explain the high rate of G:C to C:G transversions at 5′G**G**C3′+5′G**C**C3′ sites is “dNTP-stabilized misalignment” [38]. As the polymerase moves 5′ to 3′ copying the 3′C**C**G5′ triplet, the second template C could loop out, allowing a dCTP to be inserted opposite the 5′ G. If the template then realigns, a C:C mispair is created that, upon subsequent replication without correction, would produce a G:C to C:G transversion at the middle bp. Note, that the same loop-out occurring when the other strand is copied (the 3′C**G**G 5′ triplet) would not produce a mutation. Since G:C to C:G transversions at 5′G**G**C3′+5′G**C**C3′ sites are strongly biased to occur with the target C on the LGST (Figure 2f), if this event is responsible for the mutation, it must occur most often during lagging-strand synthesis (see below).

Tang *et al*. [10] pointed out that “dNTP-stabilized misalignment” is likely to be due to DNA Pol IV because it tends to make frameshift mutations that are probably also due to base loop-out. We checked to see if 5′G**G**C3′+5′G**C**C3′ sites were hotspots for G:C to C:G transversions in other strain backgrounds where mutations are less likely to be due to Pol IV. In both wild-type and MMR-defective strains it is (Figures S2f & S3f in Appendix A). Furthermore, the G:C to C:G transversions at 5′G**G**C3′+5′G**C**C3′ sites in these other strains was strongly biased to occur with the target C on the LGST, again implicating lagging-strand synthesis.

There is a strong argument against the loop-out mechanism being solely responsible for the G:C to C:G transversions at 5′G**G**C3′+5′G**C**C3′ sites. The loop-out model pairs the middle base of a triplet with the complement of the base that is 5′ to it. If this is a mismatch, a mutation will result. Of the 32 non-redundant triplet pairs, this mechanisms would fail to produce a mutation at only two (5′A**A**A3′+5′T**T**T3′ and 5′G**G**G3′+5′C**C**C3′); at eleven triplets loop-out would produce a mutation on one strand, and at eighteen triplets loop-out would produce a mutation on both strands (Table S4 in Appendix A). The BPSs that could be produced are specific to the mismatch, but from our data there is little correlation between the site-specificity of a given class of BPSs and the triplets that could produce that class by loop-out. For example, 5′C**A**A3′+5′T**T**G3′, 5′C**A**C3′+5′G**T**G3′, and 5′C**A**G3′+5′C**T**G3′ are favored sites for A:T to T:A transversions, but all three of these triplets could produce only A:T to C:G BPSs at the target base via a loop-out mechanism. Many other such examples are in the data set. Obviously there must be other factors producing site specificity.

### “Electron holes” cannot explain the site-specificity of BPSs

Spontaneously or as a result of damage, DNA bases can lose an electron, *i*.*e*. become oxidized, potentially creating a mismatch. Suárez-Villagrán *et al*. [39] found that the probability of this happening at a given base depends strongly on the adjacent bases. They calculated the probability of such an “electron hole” appearing at the middle base of each of the 64 triplets, and then compared these probabilities to published spontaneous mutation rates at these bases obtained from MA experiments with MMR-defective derivatives of four bacterial species. The correlations varied with the species and were weakest with our results for MMR-defective *E. coli* [12] (Spearman’s ρ was 0.19 with P=0.13). We checked to see if the electron hole probabilities could explain the site-specificity of the SOS-induced BPSs reported here. The only significant correlation we found was between the probabilities of electron holes at G:Cs and the rate of G:C to C:G transversions at the same triplets (Spearman’s ρ=0.55 for 5′N**G**N3′ and 0.67 for 5′N**C**N3′, P=0.03 and 0.005, respectively). However, the high rate of G:C to C:G mutations at 5′G**G**C3′+5′G**C**C3′ triplets was an outlier, as the hole probabilities at these triplets were not high enough to account for the mutation rate (*e*.*g*., 5′C**G**G3′+5′C**C**G3′ has the same hole probability as 5′G**G**C3′+5′G**C**C3′ but a G:C to C:G mutation rate 20-fold lower). We also found a weak but nonsignificant correlation between A:T to C:G transversions and 5′N**A**N3′ triplets (Spearman’s ρ = 0.34, P=0.18), but not 5′N**T**N3′ triplets (Spearman’s ρ = –0.15, P=0.59). Overall, it seems that the electron hole hypothesis cannot explain very much of the observed mutational site specificity.

### Strand-bias of the SOS-induced BPSs

In previous studies [12, 22], we reported that in MMR-defective strains there is a 2-fold bias for A:T transitions to occur with the A on the LGST rather than on the LDST, and 2-fold bias for G:C transitions to occur with C on the LGST rather than on the LDST; these results were recently confirmed for transitions in the *lacI* gene [40]. We observed weaker but similar strand biases in MMR^+^ strains [22]. In contrast to the transitions, transversions were more likely to occur with the purine on the LDST rather than on the LGST. In addition, the overall strand bias for A:T transversions was due to a few dominate sites, such as A:T to T:A mutations at 5′G**AT**C3′ sites (likely due to methylation of the A by the Dam methylase) and A:T to C:G mutations at 5′N**A**C3′ sites. At some other sites the strand bias was reversed (Figures S2 and S3 in Appendix A).

The results reported here show that the SOS-induced BPSs tend to follow these same patterns: A:T transitions are biased toward A on the LGST, G:C transitions are biased toward C on the LGST, and transversions are biased toward the purine on the LDST (Figure S1 in Appendix A). Again, local sequence context has a strong influence on these biases (Figure 2, Table S3 in Appendix A). Leaving aside sites at which no mutations with one or the other strand-bias was recovered, the range of LDST/LGST ratios varied 10- to 40- fold depending on the mutation and the site. And, as mentioned above, at some sites, the overall bias was reversed. As mentioned above, the most extreme bias was G:C to C:G transversions at 5′G**G**C3′+5′G**C**C3′ sites, which were, on average 7-fold, biased for C to be on the LGST.

### Do SOS-induced mutations occur during leading or lagging strand replication?

In the absence of overt DNA damage, DNA Pol V may gain access to the DNA when the replication fork stalls and/or skips over endogenous lesions [1]. The mutational strand bias reported here could reveal whether these events are more likely to occur during leading or lagging strand synthesis, but only if we know which mismatch is most likely to give rise to a given BPS. However, the results of *in vitro* fidelity assays, which, if on ssDNA, could identify the mismatches, are not consistent. The results of two relevant studies of Pol V fidelity, Maor-Shoshani *et al*. [41] using a gap-filling assay, and Tang *et al*. [10] using a gel-based nucleotide insertion assay, agree that for both A:T to G:C and G:C to T:A BPSs, the pyrimidine is more likely to be the template, whereas for G:C to C:G BPSs, the G is more likely to be the template. Our results show that of these three BPSs, only the G:C to C:G transversion is strongly strand-biased overall (Figure S1 in Appendix A). If G:G is the mismatch that leads to the transversion, our results predict that the G is on the leading strand template (*i*.*e*., the misinsertion of G opposite G occurs during leading strand synthesis). However, the “loop-out” mechanism discussed above predicts that C:C is the mismatch that leads to the G:C to C:G transversion, which would mean that the mismatch occurs during laggings strand synthesis. Since the mismatch giving rise to A:T to T:A transversions is strongly predicted to be T:T by the results from the Tang *et al*. (2000) study, and not strongly contradicted by the results from the Maor-Shoshani *et al*. (2000) study, from our data we can only weakly conclude that this mismatch is likely to occur during lagging strand synthesis.

Some of the SOS-induced BPSs may be due to DNA Pol IV, but the *in vitro* mismatch fidelity data for this polymerase are even more inconsistent than those for Pol V. As mentioned above, G:C to C:G transversions at 5′G**G**C3′+5′G**C**C3′ sites by loop-out could be due to Pol IV, and the C:C mismatch would take place during lagging strand synthesis (Figure 2f). But the results of three fidelity assays, two gel-based [10, 42] and one gap filling [42], are too divergent to predict which mismatches Pol IV is likely to make.

Added to these uncertainties is the fact that the strand biases we observed are strongly site-dependent (Figure 2). Gel-based fidelity assays use a template-primer pair with only one site-context for each mismatch. The single-strand gaps used in the gap-filling assays were only 350 [41] and 407 [42] Nt long, and the assays relied on production of a phenotype with the resulting restrictions on the detectable BPSs.

In conclusion, the existing fidelity data do not allow us to make strong predictions about which mismatches lead to BPSs or which DNA strand is being replicated when a given mismatch is made. Obviously, more research is needed to resolve these inconsistencies.

## Funding

This research was supported by US Army Research Office Multidisciplinary University Research Initiative (MURI) Award [W911NF-09-1-0444 to P.L.F. & H.T.] and the National Institutes of Health [T32 GM007757 to B.A.N]. The funding sources had no role in the study design; in the collection, analysis and interpretation of data; in the writing of the report; and in the decision to submit the article for publication.

## Author contributions

Brittany A. Niccum: Conceptualization, Methodology, Formal analysis, Investigation, Writing - Original Draft, Supervision. Christopher P. Coplen: Methodology, Investigation, Writing - Review & Editing. Heewook Lee: Methodology, Software, Formal analysis, Data Curation, Writing - Review & Editing. Wazim MohammedIsmail: Methodology, Software, Formal analysis, Data Curation. Haixu Tang: Conceptualization, Methodology, Software, Formal analysis, Data Curation, Writing - Review & Editing, Supervision, Project administration, Funding acquisition. Patricia L. Foster: Conceptualization, Methodology, Formal analysis, Writing - Review & Editing, Supervision, Project administration, Funding acquisition.

## Declaration of Competing Interests

Brittany A. Niccum is currently employed at a life-sciences company but was not so employed when the research was done and analyzed.

## Acknowledgements

We thank the following past members of the P.L.F. laboratory for technical assistance: H. Bedwell, J. Eagan, N. Gruenhagen, E. Popodi, I. Rameses, D. Simon, and J. Townes. We are especially grateful to R. Woodgate and J.P. McDonald, NIH-NICHD, for performing Western blots for us. The National BioResource Project at the (Japanese) National Institute of Genetics provided bacterial strains and plasmids. This research was supported by US Army Research Office Multidisciplinary University Research Initiative (MURI) Award [W911NF-09-1-0444 to P.L.F. & H.T.] and the National Institutes of Health [T32 GM007757 to B.A.N].

