## Supplemental Materials for "New complexities of SOS-induced “untargeted” mutagenesis in *Escherichia coli* as revealed by mutation accumulation and whole-genome sequencing"

**Table S1. Bacterial strains**

| Strain | Relevant genotype | Donor | Recipient | Target | Method | Reference |
|---|---|---|---|---|---|---|
| PFM2 | MG1665 *rph*$^+$ | | | | | [1] |
| PFM20 | Δ*sulA* | JW0941 | | *sulA* | A | [2] |
| PFM50 | PFM2 Δ*umuDC* | | | | | [3] |
| PFM117 | Δ*sulA*/pKD46 | Plasmid DNA | PFM20 | λ red recombinase | B | [4] |
| PFM128 | Δ*recA*-*cat*-I-SceI Δ*sulA* | PCR product | PFM117 | *recA* | C | [5] |
| PFM129 | *recA730* Δ*sulA* | PC1427 | PFM20 | Δ*recA*-*cat*-I-SceI | D | [6] |
| PFM138 | *recA730* Δ*metA*::Kn$^R$ Δ*sulA* | JW3973 | PFM129 | *metA* | E | [2] |
| **lexA*-Kn$^R$ cassette | *lexA*-Kn$^R$ Δ*sulA* | PCR product | PFM117 | *lexA* | F | This Study |
| PFM196 | *recA730* Δ*lexA* Δ*sulA* | *lexA*-Kn$^R$ cassette | PFM129 | *lexA* | A | This Study |
| PFM199 | *recA730 lexA3* Δ*sulA* | C231 | PFM138 | *lexA* Δ*metA*::Kn$^R$ | G | [7] |
| PFM520 | Δ*sulA*Δ | JW0941 | PFM50 | *sulA* | A | [2] |
| PFM588 | Δ*recA*-*cat*-I-SceI Δ*sulA* Δ*umuDC* | PFM128 | PFM50 | *recA* | H | |
| PFM623 | *recA730* Δ*sulA* Δ*umuDC* | PFM129 | PFM520 | Δ*recA*-*cat*-I-SceI | I | This Study |
| PFM646 | *recA730* Δ*lexA* Δ*sulA* Δ*umuDC* | *lexA*-Kn$^R$ cassette | PFM623 | lex*A* | A | This Study |
| PFM673 | *recA730* Δ*lexA* Δ*sulA* Δ*umuDC* Δ*dinB* | JW0221 | PFM646 | *dinB* | A | [2] |

A. P1 phage transduction of a $Kn^R$ deletion-insertion in the target gene, or $Kn^R$ linked to the desired allele of the target gene, selecting for $Kn^R$, followed by FLP recombination to remove the $Kn^R$ element, and in the case of *lexA*, most of the gene [4]. B. Transformation with plasmid DNA, selecting for $Carb^R$. C. Creation of a Δ*recA*-*cat*-I-SceI cassette followed by recombination onto the chromosome [5]. D. Transformation with a PCR product of the *recA730* mutant gene selecting for resistance to anhydrotetracycline to replace the Δ*recA*-*cat*-I-SceI cassette. E. P1 phage transduction selecting for $Kn^R$. F. Creation of $Kn^R$ cassette linked to the desired allele followed by recombination into the genome [4]. G. P1 phage transduction selecting for growth on minimal medium to remove the $Kn^R$ element, replace the auxotrophy, and transfer the linked *lexA3* allele. H. P1 phage transduction selecting for $Cm^R$. I. P1 phage transduction selecting for resistance to anhydrotetracycline to replace the Δ*recA*-*cat*-I-SceI cassette with *recA730*. *The strain carrying this cassette, which was the donor for P1 transductions, was not saved.

**Table S2. Parameters of the MA experiments**

| Strain | Description | No. of MA lines | No. of generations | No. of BPSs | *BPSs per generation x $10^3$ | *95% CL x $10^3$ | No. of Indels | *Indels per generation x $10^3$ | *95% CL x $10^3$ |
|---|---|---|---|---|---|---|---|---|---|
| #Collective | Wild type | 342 | 2015066 | 1933 | 0.96 | 0.11 | 193 | 0.097 | 0.018 |
| PFM129 | *recA730 ΔsulA* | 40 | 21921 | 621 | 28.8 | 2.5 | 28 | 1.3 | 0.4 |
| PFM196 | *recA730 ΔlexA ΔsulA* | 38 | 19699 | 804 | 40.8 | 2.9 | 16 | 0.81 | 0.54 |
| PFM673 | *recA730 ΔlexA ΔsulA ΔumuDC ΔdinB* | 47 | 52138 | 67 | 1.29 | 0.35 | 9 | 0.17 | 0.11 |
| PFM199 | *recA730 lexA3 ΔsulA* | 40 | 102339 | 146 | 1.43 | 0.27 | 15 | 0.15 | 0.09 |

*Data are means ± 95% CL. Statistical calculations were as described [8]. #The combined results of eight experiments with strains with wild-type mutational phenotypes [8]. CL, confidence limit.

### Table S3. Statistical evaluation of the strand and site specificity of BPS rates

| | BPSs/Generation/Nt X $10^{10}$ | | | | | | |
|---|---|---|---|---|---|---|---|
| | Purine on the LGST | | Purine on the LDST | | | | |
| 5' 3'+5' 3' | Mean | SD | Mean | SD | LGST-LDST | *t | P |
| | | | **A:T to G:C** | | | | |
| AAA+TTT | 6.6 | 0.30 | 22 | 0.37 | -15 | 46 | 0.01 |
| AAC+GTT | 20 | 0.46 | <3 | >0.35 | >17 | 42 | 0.02 |
| AAG+CTT | 7.6 | 0.83 | 15 | 0.78 | -7.5 | 9.4 | 0.07 |
| AAT+ATT | 2.9 | 0.32 | 2.9 | 0.35 | 0.01 | 0.0 | 0.97 |
| CAA+TTG | 6.3 | 0.68 | 9.4 | 0.42 | -3.1 | 5.5 | 0.12 |
| CAC+GTG | 18 | 1.4 | <4 | >0.44 | >14 | 14 | 0.05 |
| CAG+CTG | 11 | 0.19 | 7.0 | 0.23 | 4.5 | 21 | 0.03 |
| CAT+ATG | 9.4 | 1.0 | 3.2 | 0.34 | 6.2 | 8.1 | 0.08 |
| GAA+TTC | 46 | 0.95 | 11 | 0.59 | 35 | 44 | 0.01 |
| GAC+GTC | 18 | 1.1 | 13 | 0.43 | 4.3 | 5.0 | 0.13 |
| GAG+CTC | 5.6 | 0.61 | 5.7 | 0.62 | -0.04 | 0.1 | 0.96 |
| GAT+ATC | 25 | 1.8 | 17 | 0.10 | 8.3 | 6.7 | 0.09 |
| TAA+TTA | 3.5 | 0.38 | 14 | 0.89 | -10 | 15 | 0.04 |
| TAC+GTA | 9.1 | 1.00 | 18 | 0.11 | -9.2 | 13 | 0.05 |
| TAG+CTA | 18 | 2.0 | <8 | >1.1 | >10 | 6.3 | 0.10 |
| TAT+ATA | 11 | 0.50 | 23 | 1.0 | -11 | 14 | 0.04 |
| All A:T | 14 | 0.09 | 11 | 0.03 | 3.3 | 52 | 0.01 |
| | | | **G:C to A:T** | | | | |
| AGA+TCT | 4.3 | 0.47 | 38 | 2.7 | -34 | 18 | 0.04 |
| AGC+GCT | 15 | 0.25 | 18 | 0.80 | -2.9 | 4.9 | 0.13 |
| AGG+CCT | 14 | 0.64 | 19 | 0.12 | -4.7 | 10 | 0.06 |
| AGT+ACT | 9.7 | 0.06 | 14 | 0.47 | -4.8 | 14 | 0.04 |
| CGA+TCG | 3.3 | 0.37 | 30 | 0.20 | -27 | 92 | 0.01 |
| CGC+GCG | 29 | 0.18 | 44 | 0.93 | -14 | 21 | 0.03 |
| CGG+CCG | 14 | 0.40 | 30 | 0.13 | -17 | 55 | 0.01 |
| CGT+ACG | 9.9 | 0.32 | 23 | 0.52 | -13 | 30 | 0.02 |
| GGA+TCC | 4.3 | 0.52 | 4.3 | 0.52 | 0.0 | 0.0 | 0.98 |
| GGC+GCC | 10 | 0.67 | 18 | 0.78 | -7.7 | 11 | 0.06 |
| GGG+CCC | 5.1 | 0.61 | 10 | 1.2 | -5.0 | 5.2 | 0.12 |
| GGT+ACC | 19 | 0.87 | 6.4 | 0.04 | 13 | 21 | 0.03 |
| TGA+TCA | 12 | 1.4 | 17 | 1.2 | -5.6 | 4.3 | 0.15 |
| TGC+GCA | 10 | 0.64 | 48 | 0.58 | -37 | 61 | 0.01 |
| TGG+CCA | 8.5 | 0.27 | 11 | 0.71 | -2.7 | 4.9 | 0.13 |
| TGT+ACA | 25 | 1.7 | 20 | 0.60 | 4.2 | 3.2 | 0.19 |
| All G:C | 13 | 0.20 | 24 | 0.15 | -11 | 62 | 0.01 |

| | | | | | | | |
|---|---|---|---|---|---|---|---|
| | | | **A:T to T:A** | | | | |
| AAA+TTT | 15 | 0.16 | 20 | 0.88 | -4.2 | 6.7 | 0.09 |
| AAC+GTT | 12 | 1.4 | 12 | 0.74 | 0.0 | 0.0 | 1.00 |
| AAG+CTT | 3.8 | 0.41 | 7.5 | 0.82 | -3.8 | 5.8 | 0.11 |
| AAT+ATT | 12 | 1.3 | 17 | 0.11 | -5.7 | 6.4 | 0.10 |
| CAA+TTG | 34 | 0.87 | 103 | 3.2 | -69 | 30 | 0.02 |
| CAC+GTG | 22 | 0.13 | 164 | 1.4 | -142 | 140 | 0.005 |
| CAG+CTG | 28 | 0.36 | 180 | 0.30 | -152 | 460 | 0.001 |
| CAT+ATG | 3.1 | 0.34 | 22 | 2.0 | -19 | 14 | 0.05 |
| GAA+TTC | 17 | 0.56 | 34 | 3.5 | -17 | 6.8 | 0.09 |
| GAC+GTC | 8.8 | 1.1 | 9 | 1.1 | -0.1 | 0.1 | 0.95 |
| GAG+CTC | 22 | 1.2 | 34 | 0.21 | -11 | 14 | 0.05 |
| GAT+ATC | 11 | 0.07 | 19 | 0.84 | -8.3 | 14 | 0.05 |
| TAA+TTA | 31 | 0.21 | 94 | 1.8 | -63 | 49 | 0.01 |
| TAC+GTA | 9.1 | 0.06 | 37 | 0.23 | -27 | 167 | 0.004 |
| TAG+CTA | 9.0 | 0.98 | 79 | 1.5 | -70 | 55 | 0.01 |
| TAT+ATA | 23 | 0.73 | 19 | 1.4 | 3.7 | 3.2 | 0.19 |
| All A:T | 17 | 0.24 | 55 | 1.0 | -38 | 50 | 0.01 |
| | | | **G:C to T:A** | | | | |
| AGA+TCT | 17 | 0.11 | 34 | 0.21 | -17 | 100 | 0.01 |
| AGC+GCT | 3.0 | 0.33 | 5.9 | 0.04 | -3.0 | 13 | 0.05 |
| AGG+CCT | 24 | 0.70 | 9.5 | 1.0 | 14 | 16 | 0.04 |
| AGT+ACT | <5 | >0.53 | <5 | >0.53 | 0.0 | UD | UD |
| CGA+TCG | 20 | 0.65 | 95 | 1.4 | -75 | 70 | 0.01 |
| CGC+GCG | 8.4 | 0.54 | 29 | 0.30 | -21 | 48 | 0.01 |
| CGG+CCG | 14 | 0.40 | 52 | 2.6 | -39 | 21 | 0.03 |
| CGT+ACG | 6.6 | 0.80 | 36 | 0.60 | -30 | 42 | 0.02 |
| GGA+TCC | 17 | 1.1 | 8.6 | 0.05 | 8.6 | 11 | 0.06 |
| GGC+GCC | 13 | 0.38 | 5.2 | 0.56 | 7.9 | 16 | 0.04 |
| GGG+CCC | 46 | 0.30 | 10 | 0.06 | 35 | 162 | 0.004 |
| GGT+ACC | 26 | 1.7 | <3 | >0.39 | 23 | 19 | 0.03 |
| TGA+TCA | 35 | 2.9 | 83 | 0.48 | -48 | 24 | 0.03 |
| TGC+GCA | 10 | 0.64 | 15 | 0.48 | -4.9 | 8.6 | 0.07 |
| TGG+CCA | 48 | 2.6 | 14 | 0.24 | 34 | 18 | 0.03 |
| TGT+ACA | 16 | 0.10 | 20 | 1.3 | -4 | 4.4 | 0.14 |
| All G:C | 18 | 0.35 | 27 | 0.05 | -9.0 | 36 | 0.02 |
| | | | **A:T to C:G** | | | | |
| AAA+TTT | 4.4 | 0.03 | 44 | 0.27 | -39 | 205 | 0.003 |
| AAC+GTT | 12 | 0.60 | 20 | 0.46 | -8.7 | 16 | 0.04 |
| AAG+CTT | 38 | 0.23 | 87 | 4.5 | -49 | 16 | 0.04 |
| AAT+ATT | 2.9 | 0.35 | 12 | 0.59 | -8.6 | 18 | 0.04 |

| | | | | | | | |
|---|---|---|---|---|---|---|---|
| CAA+TTG | 19 | 0.61 | 31 | 0.91 | -12 | 16 | 0.04 |
| CAC+GTG | 3.6 | 0.44 | 3.6 | 0.40 | 0.0 | 0.1 | 0.95 |
| CAG+CTG | 21 | 0.92 | 54 | 0.06 | -33 | 51 | 0.01 |
| CAT+ATG | 9.4 | 0.42 | 13 | 0.08 | -3.2 | 11 | 0.06 |
| GAA+TTC | 12 | 0.59 | 2.9 | 0.35 | 8.6 | 18 | 0.04 |
| GAC+GTC | 8.8 | 0.05 | 18 | 2.2 | -8.9 | 5.9 | 0.11 |
| GAG+CTC | 11 | 1.4 | 68 | 1.7 | -57 | 36 | 0.02 |
| GAT+ATC | 25 | 1.1 | 28 | 0.81 | -2.8 | 2.8 | 0.22 |
| TAA+TTA | 14 | 0.9 | 66 | 3.2 | -52 | 22 | 0.03 |
| TAC+GTA | <5 | >0.55 | 4.6 | 0.56 | >0 | 0.8 | 0.58 |
| TAG+CTA | <9 | >1.1 | 79 | 5.6 | -70 | 18 | 0.04 |
| TAT+ATA | 7.5 | 0.05 | 7.6 | 0.05 | 0.0 | 1.0 | 0.49 |
| All A:T | 12 | 0.2 | 32 | 0.92 | -19 | 28 | 0.02 |
| **G:C to C:G** | | | | | | | |
| AGA+TCT | 8.7 | 1.1 | 13 | 1.5 | -4.1 | 3.1 | 0.20 |
| AGC+GCT | 0.0 | 0.00 | 39 | 1.3 | -39 | 43 | 0.01 |
| AGG+CCT | 4.8 | 0.58 | 14 | 0.64 | -9.5 | 16 | 0.04 |
| AGT+ACT | <5 | >0.53 | 14 | 0.47 | >9 | 19 | 0.03 |
| CGA+TCG | <3 | >0.37 | 14 | 0.70 | >11 | 19 | 0.03 |
| CGC+GCG | 8.4 | 0.05 | 50 | 1.3 | -41 | 46 | 0.01 |
| CGG+CCG | 17 | 1.4 | 11 | 0.07 | 5.6 | 5.7 | 0.11 |
| CGT+ACG | 9.9 | 0.44 | 16 | 0.28 | -6.5 | 18 | 0.04 |
| GGA+TCC | 8.6 | 0.94 | 17 | 0.11 | -8.5 | 13 | 0.05 |
| GGC+GCC | 70 | 0.13 | 504 | 30 | -434 | 20 | 0.03 |
| GGG+CCC | 5.1 | 0.61 | 45 | 3.2 | -40 | 18 | 0.04 |
| GGT+ACC | 13 | 0.67 | 51 | 1.1 | -38 | 43 | 0.01 |
| TGA+TCA | 14 | 0.24 | 8.6 | 0.38 | 5.8 | 18 | 0.04 |
| TGC+GCA | 7.6 | 0.34 | 23 | 0.43 | -15 | 39 | 0.02 |
| TGG+CCA | 11 | 0.72 | 19 | 0.20 | -8.2 | 15 | 0.04 |
| TGT+ACA | 4.1 | 0.50 | 12 | 0.55 | -8.2 | 16 | 0.04 |
| All G:C | 13 | 0.24 | 62 | 2.9 | -49 | 24 | 0.03 |
| All G:C without GGC+CGG | 8.0 | 0.25 | 24 | 0.56 | -16 | 38 | 0.02 |

Numbers are the means and 95% CL of the combined conditional BPSs rates of the *recA730* (PFM129) and the *recA730* Δ*lexA* (PFM196) mutant strains; both strains were also Δ*sulA*. Statistical calculations were as described [8]. Triplets are written 5′ to 3′ with the target base in the center. Because of the small numbers, the difference between the results for LGST and LDST was a more stable statistic than the ratio; *t, Student's t; t and the probability were calculated as described [9]. When a category had 0 BPS, t and P were calculated using the least detectable number of BPS. UD, undefined; CL, confidence limits.

**Table S4: Comparison of the BPSs predicted by dNTP-stabilized misalignment and the SOS-induced BPS rates observed**

| Triplet | Predicted BPS at center base-pair | | BPS/Generation/Nt x $10^{10}$ | | | | | |
|---|---|---|---|---|---|---|---|---|
| | | | Mean | 95% CL | Mean | 95% CL | Mean | 95% CL |
| 5' 3'+5 '3' | | | A:T>G:C | | A:T>T:A | | A:T>C:G | |
| AAA+TTT | NONE | NONE | 14 | 0.1 | 18 | 0.7 | 24 | 0.3 |
| AAC+GTT | NONE | AT>CG | 10 | 0.5 | 12 | 2.1 | 16 | 0.1 |
| AAG+CTT | NONE | AT>GC | 11 | 1.6 | 5.7 | 1.2 | 62 | 4.7 |
| AAT+ATT | NONE | AT>TA | 2.9 | 0.04 | 14 | 1.1 | 7.2 | 0.2 |
| CAA+TTG | AT>CG | NONE | 7.8 | 0.3 | 69 | 2.3 | 25 | 0.3 |
| CAC+GTG | AT>CG | AT>CG | 9.0 | 1.4 | 92 | 1.5 | 3.6 | 0.0 |
| CAG+CTG | AT>CG | AT>GC | 9.3 | 0.4 | 103 | 0.1 | 37 | 1.0 |
| CAT+ATG | AT>TA | AT>TA | 6.3 | 1.4 | 13 | 1.6 | 11 | 0.5 |
| GAA+TTC | AT>GC | NONE | 29 | 0.4 | 26 | 3.0 | 7.2 | 0.2 |
| GAC+GTC | AT>GC | AT>CG | 15 | 0.7 | 8.8 | 2.1 | 13 | 2.2 |
| GAG+CTC | AT>GC | AT>GC | 5.6 | 1.2 | 28 | 0.9 | 39 | 3.1 |
| GAT+ATC | AT>TA | AT>GC | 21 | 1.9 | 15 | 0.8 | 26 | 1.9 |
| TAA+TTA | AT>TA | NONE | 8.7 | 0.5 | 63 | 1.6 | 40 | 4.1 |
| TAC+GTA | AT>CG | AT>TA | 14 | 0.9 | 23 | 0.3 | 2.3 | 0.6 |
| TAG+CTA | AT>GC | AT>TA | 8.9 | 1.9 | 44 | 0.5 | 40 | 5.6 |
| TAT+ATA | AT>TA | AT>TA | 17 | 1.5 | 21 | 0.7 | 7.6 | 0.1 |
| | | | G:C>A:T | | G:C>C:G | | G:C>T:A | |
| AGA+TCT | GC>AT | GC>AT | 21 | 2.2 | 11 | 2.6 | 26 | 0.3 |
| AGC+GCT | GC>AT | GC>CG | 16 | 0.5 | 19 | 1.3 | 4.5 | 0.3 |
| AGG+CCT | GC>AT | NONE | 17 | 0.8 | 9.5 | 1.2 | 17 | 0.3 |
| AGT+ACT | GC>TA | GC>AT | 12 | 0.4 | 7.2 | 0.5 | 0 | 0 |
| CGA+TCG | GC>CG | GC>AT | 17 | 0.6 | 6.7 | 0.7 | 57 | 0.7 |
| CGC+GCG | GC>CG | GC>CG | 37 | 0.8 | 29 | 1.3 | 19 | 0.2 |
| CGG+CCG | NONE | GC>CG | 22 | 0.3 | 14 | 1.4 | 33 | 2.9 |
| CGT+ACG | GC>TA | GC>CG | 16 | 0.2 | 13 | 0.2 | 21 | 1.4 |
| GGA+TCC | NONE | GC>AT | 4.3 | 1.0 | 13 | 0.8 | 13 | 1.1 |
| GGC+GCC | GC>CG | NONE | 14 | 0.1 | 288 | 30 | 9.1 | 0.2 |
| GGG+CCC | NONE | NONE | 7.6 | 1.8 | 25 | 3.8 | 28 | 0.2 |
| GGT+ACC | GC>TA | NONE | 13 | 0.9 | 32 | 0.4 | 13 | 1.6 |
| TGA+TCA | GC>AT | GC>TA | 14 | 0.2 | 11 | 0.1 | 59 | 2.4 |
| TGC+GCA | GC>CG | GC>TA | 29 | 1.2 | 15 | 0.8 | 13 | 0.2 |
| TGG+CCA | NONE | GC>TA | 9.8 | 0.4 | 15 | 0.5 | 31 | 2.8 |
| TGT+ACA | GC>TA | GC>TA | 23 | 1.1 | 8.2 | 1.0 | 18 | 1.2 |

Numbers are the means and 95% CL computed using the combined BPSs/generation/Nt results of the *recA730* (PFM129) and *recA730* ∆*lexA* (PFM196) mutant strains, both of which are also ∆*sulA.* Statistical calculations were as described [8]. Triplets are written 5′ to 3′ with the target base in the center. CL, confidence limits.

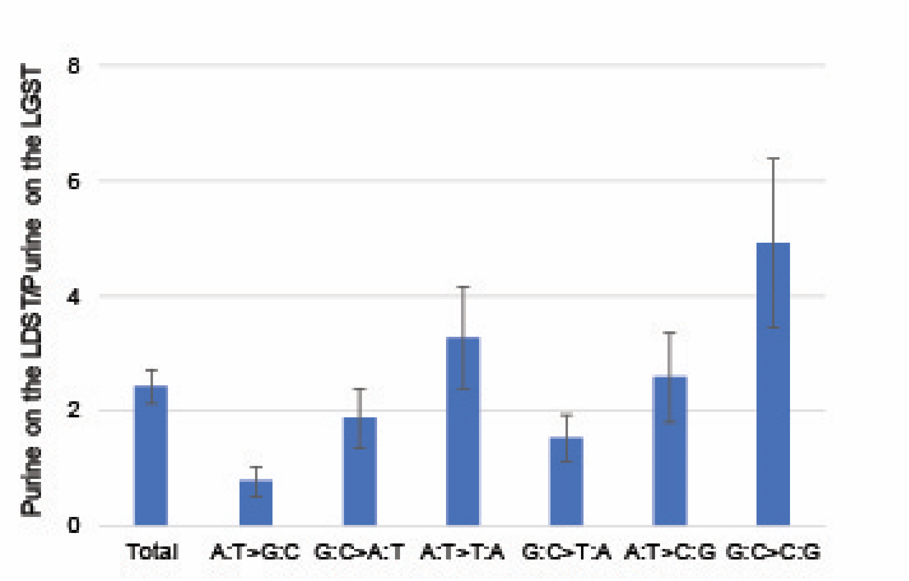


Figure S1: The ratio of the rates of SOS-induced base-pair substitutions when the purine is on the LDST and the pyrimidine on the LGST versus when in the opposite orientation. The ratios were computed using the combined BPSs/generation results of the *recA730* (PFM129) and *recA730* Δ*lexA* (PFM196) mutant strains, both of which are also Δ*sulA*. LGST = DNA lagging strand template; LDST = DNA leading strand template. Bars represent means and the error bars are 95% CLs, calculated as described [8, 10]. CL, confidence limits.

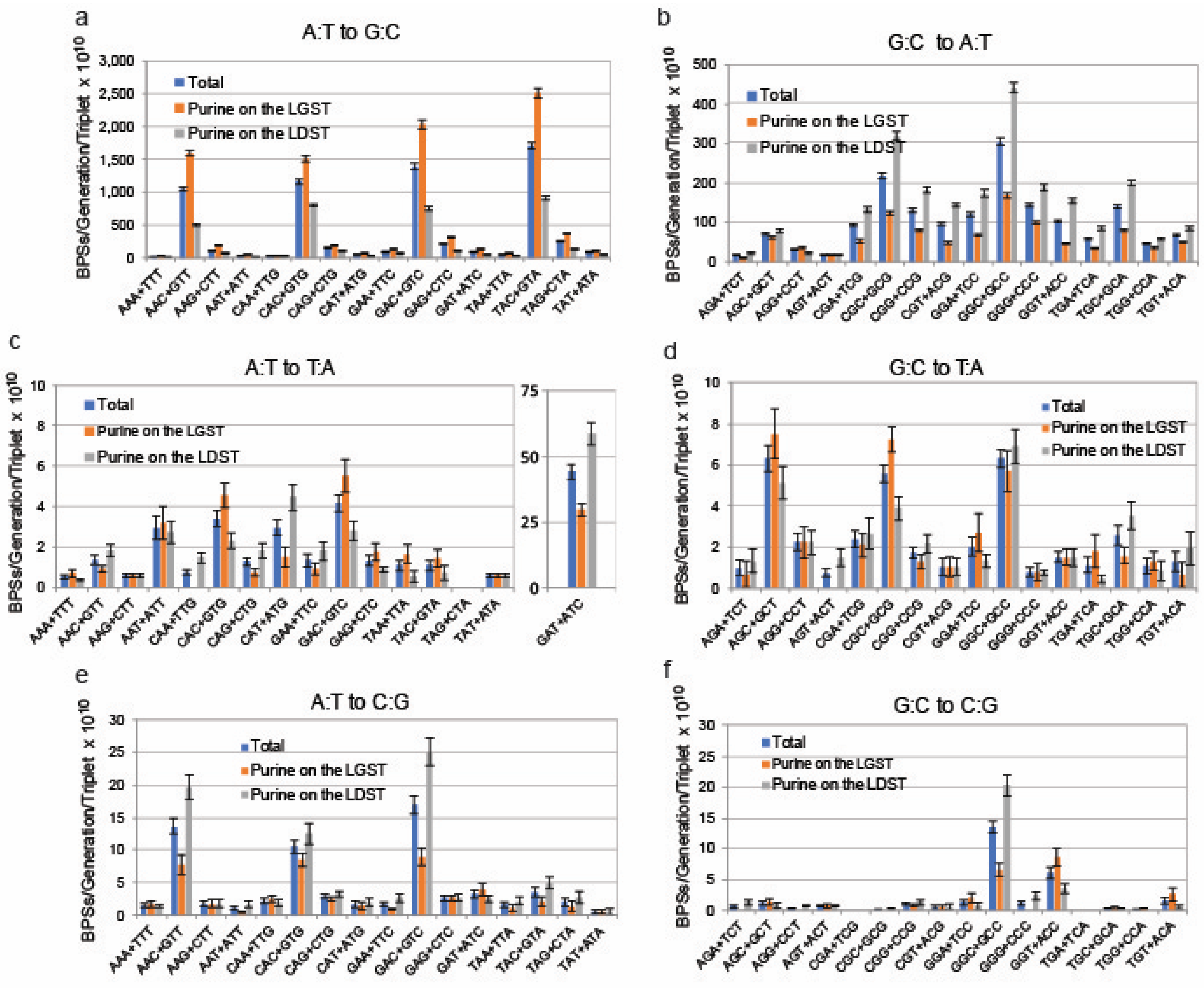


Figure S2: The site and strand specificity of base-pair substitutions from strains defective for mismatch repair. Conditional mutation rates were computed from the combined results of ten experiments with MMR-defective strains [8]. LGST = DNA lagging strand template; LDST = DNA leading strand template. Triplets are written 5′ to 3′ with the target base in the center. Bars represent means and the error bars are 95% CLs computed as described [8]. Note change of scales among the charts and the extra chart in Fig. S2c for results at 5′GAT3′ + 5′ATC3′ sites. CL, confidence limits.

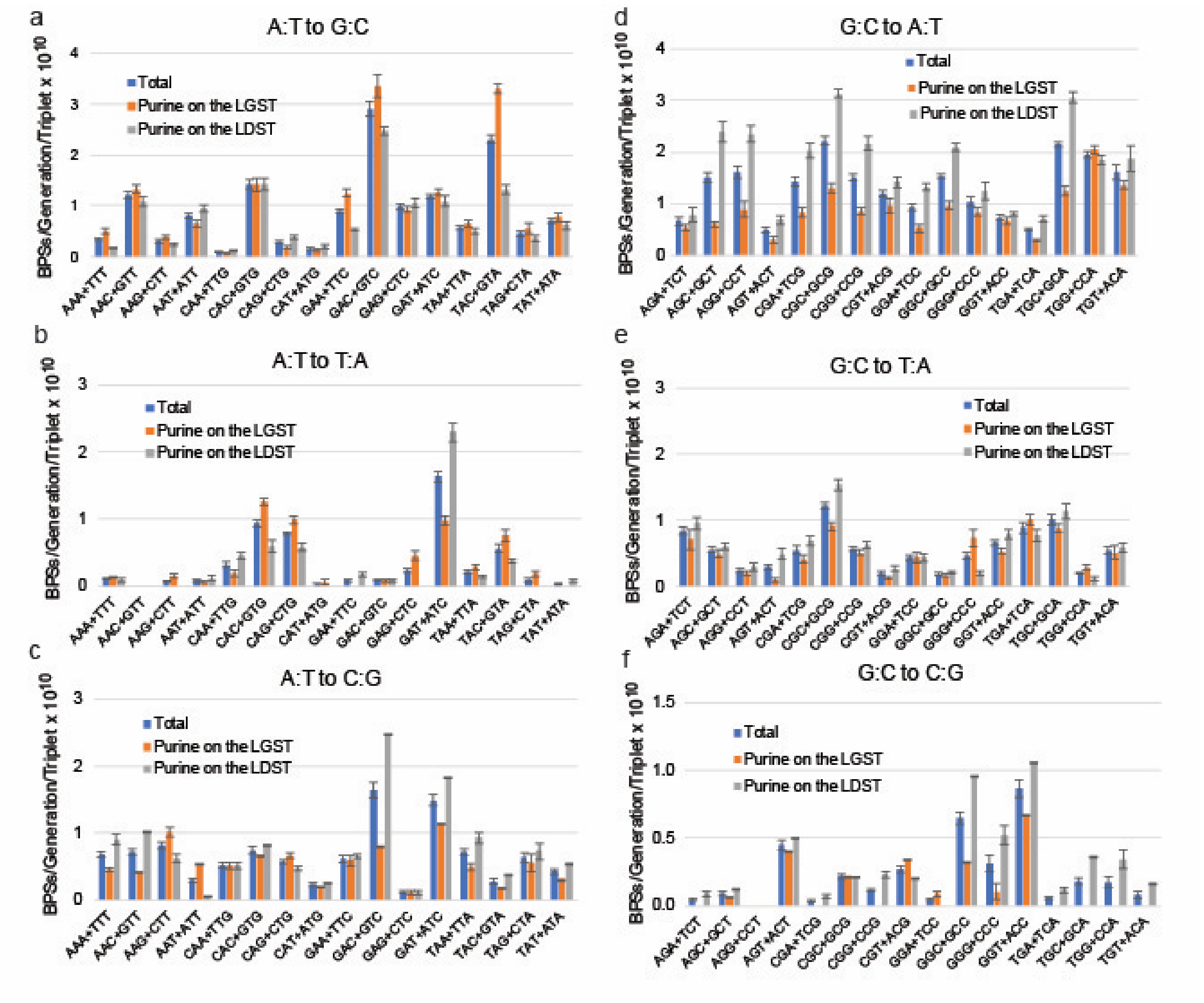


Figure S3: The site and strand specificity of base-pair substitutions from wild-type strains. Conditional mutation rates were computed from the combined results of eight experiments with strains with wild-type mutational phenotypes [8]. LGST = DNA lagging strand template; LDST = DNA leading strand template. Triplets are written 5′ to 3′ with the target base in the center. Bars represent means and the error bars are 95% CLs computed as described [8]. Note change of scales among the charts. CL, confidence limits.
